# A 6-week all-liquid caloric restriction diet reduced abundance of some butyrate-producing bacteria while increasing abundance of mucin-degrading bacteria

**DOI:** 10.64898/2026.11.23.768597

**Authors:** Blake Dirks, Khemlal Nirmalkar, Susanne Votruba, Tim Hollstein, Jonathan Krakoff, Rosa Krajmalnik-Brown

## Abstract

Caloric restriction diets are a common diet strategy to reduce excess weight and obesity. The diets work by lowering the number of calories or energy input below a human’s normal energy expenditure creating an energy deficit. Caloric restriction diet meals can be composed of whole foods, meal replacement shakes, or a combination of the two. While the effects of these diets on human metabolism are well-known, the effects of these diets and the durability of these effects on the human gut microbiome are less well-known. The objective of this work was to investigate the effects of a 6-week all-liquid caloric restriction diet on the human gut microbiota using 16S rRNA gene amplicon sequencing to assess changes in diversity, community composition, and predicted function. We found that during the diet period beta-diversity shifts significantly away from what it was before the intervention (baseline), and returns to baseline a week after the diet. During the diet period we also observed a decrease in abundance of fiber-degrading and butyrate-producing bacteria with a simultaneous increase in abundance of mucin-degrading bacteria with a return to baseline level abundances in the post-diet period. Despite a liquid diet relatively high in carbohydrate (59% carbohydrate, 25% fat, 16% protein), predicted function analysis showed a shift in substrate preference away from dietary carbohydrates towards protein and host-glycans during the diet period. These findings suggest that despite the overall high carbohydrate content all-liquid caloric restriction diets deprive the gut microbiota of their preferred nutrient source due to the overall decline in calorie delivery which forces them to consume alternative nutrient sources like protein and host-glycans.

## Introduction

Caloric restriction (CR) is a well-known dietary intervention to reduce obesity^1^. CR diet regimes ideally maintain a balanced macronutrient composition while reducing total energy intake putting an individual at an energy deficit but avoiding malnutrition^2^. Researchers have found that CR diets reduced adiposity^3^, improved insulin-sensitivity, reduced risk factors for cardiovascular disease^4^, and increased lifespans in many mammals, fish, and even yeast^5,6^. Although the human lifespan makes similar observations very difficult, studies have found evidence that CR alters metabolic pathways associated with longevity^7,8^

Because diet is one of the main drivers that shape the gut microbiome^9,10^ researchers have investigated how CR diets influence the gut microbiome^11–13^. These CR diet studies showed alterations in beneficial members of the gut microbiota, for example *Ruminococcus faecis* and *Bifidobacterium spp.*, important fiber degraders^11^, and *Akkermansia spp.*, an important mucin-degrading bacterium^14^, increased. Additionally, the abundances of Akkermansia and Ruminococcaceae have been positively associated with improvements in blood pressure and weight-loss^15^ healthy fasting glucose and triglycerides and fat distribution^16^.

The CR diet in many of the human studies consisted of either whole foods or a combination of whole foods and liquid diets with a moderate caloric reduction of 10% – 30% of daily caloric intake^17^Some CR diets though, such as the Very Low Energy Diet (VLED) or Very Low Caloric Energy Diet (VLCD), are all-liquid diets that provide around 800kcal/day^18^. These diets are designed to be readily absorbable by the small intestine with very few nutrients reaching the colon^19^, and are prescribed for patients in limited cases, for example extreme obesity or in preparation for surgical procedures such as bariatric surgery, and under physician observation to prevent nutrient deficiencies^20^. These types of diets result in significant weight loss and improvement of metabolic outcomes such as reduced insulin resistance, LDL and HDL levels, and improved inflammatory measures^21^. However, many popular full-liquid meal replacements CR diet products are available to the public, such as Ensure and Slim Fast, without the benefit of physician observation. The effects of long-term use of all-liquid CR diets on the gut microbiome have not been well studied, thus the possible risks and benefits of these diets in the context of the microbiome remain generally unknown.

In this study, we analyzed changes in microbial community structure and predicted function from 16S rRNA gene amplicon sequencing over the course of a 6-week all-liquid CR diet including the pre-diet baseline and post-diet period. We hypothesized that the all-liquid CR diet would be readily absorbed by the human host and reduce the amount of nutrients reaching the colon compared to the weight maintenance diet periods. The lack of nutrition in the form of carbohydrates and protein reaching the colon in the all-liquid CR diets most likely would enrich bacteria that feed on host-derived carbohydrates and deplete fiber degrading bacteria and bacteria associated with those fiber degraders, such as butyrate producers. This trend would be opposite trends recorded in whole-food CR diets, where fiber degraders and associated butyrate producers increased^22^.

## Methods and Materials

### Study Overview

16S rRNA gene amplicon sequences for this manuscript were generated from a previously published study focusing on weight loss in obese individuals during a CF diet by the National Institute of Diabetes and Digestive and Kidney Diseases (NIDDK) in Phoenix, AZ^23^.

For this study,47 obese individuals were screened, 15 were admitted based on health examinations, laboratory analysis, electrocardiograph, and physiological assessment, and three individuals did not complete the study. 12 participants were admitted for 77 ± 4 days to an inpatient unit of the Obesity and Diabetes Clinical Research Section of the Phoenix, AZ NIDDK. Recruitment was conducted in the general Phoenix metropolitan area, with no direct recruitment from Phoenix Indian Medical Center or any Indian Health Service facility.

Participant characteristics are given in **Supplementary Table 1**. Participants were fed a weight-maintaining diet composed of 50% carbohydrate, 30% fat, and 20% protein for 21 days to establish a baseline metabolic level. Caloric intake was adjusted to maintain a stable body weight.

Next, participants were put on an all-liquid caloric restriction diet of Ensure, containing a macronutrient composition of 59% carbohydrate, 25% fat, 16% protein, that provided 50% of the calories needed to maintain their weight calculated from the baseline period. Participants consumed this diet for 42 days. Participants were then placed on a post-caloric restriction diet for 14 days, where they consumed a weight-maintaining diet similar to the baseline period.

### Fecal Sample Collection, Bomb Calorimetry, DNA Extraction, and Sequencing

Fecal samples were collected three consecutive days per week. Fecal samples for bomb calorimetry were stored at –20C until use. Fecal samples for sequencing were stored at –80C until use.

Methods for sample processing for bomb calorimetry are detailed in Reinhardt, et al., 2015.^24^ Bomb calorimetry was carried out using the Parr Isoperibol Calorimeter 6200 with a model 1108 oxygen bomb according to manufacterer’s protocol. Samples were processed on ice and extracted using the Qiagen Powersoil DNA extraction kit. Library preparation was performed according to the Earth Microbiome Project protocol (https://earthmicrobiome.org/protocols-and-standards/16s/). Extracted DNA was sequenced on an Illumina MiSeq with 2X300 bp chemistry using 515F-806R primer set^25^, targeting the V4 region of the 16S rRNA gene.

### Sequencing Data Processing

Sequencing data were processed and analyzed with the Qiime2 pipeline^26^. Sequence quality control was performed with DADA2 (Callahan et al., 2016). α-diversity analysis was calculated with observed ASVs, Pielou’s Evenness, Shannon Diversity, and Inverse Simpson Index. β-diversity analysis was calculated with weighted and unweighted Unifrac^27^ (Supplementary Figure 1), Bray-Curtis dissimilarity^28^, and Jaccard similarity^29^ metrics. Taxonomic analysis was performed using the Phyloseq R Package^30^. Functional pathway and gene prediction was performed with PICRUSt2^31^. All figures were created using the ggplot2 R Package^32^.

### Differential Abundance Analysis

Differential abundances of predicted functional pathways and genes were calculated using the R program Masslin2^33^. Pathway and gene features were normalized by total sum scaling (TSS) where counts for each feature were divided by the total counts in a sample. Features with a count of zero were given a small pseudo-value (half of the lowest non-zero value), then all features were log-transformed. Differential abundance results were produced by a linear model comparing abundances at each period to abundances at baseline. P values were corrected for multiple comparisons using the Benjamini-Hochberg (BH) method. P values < 0.05 were considered statistically significant. Adjusted P values < 0.05 were considered statistically significant.

## Results and Discussion

### Microbial diversity was altered by the CR diet, but returned to baseline by the post-diet period

Because diet is one of the strongest drivers of change in the gut microbiome^9,34^, we assessed fecal energy concentration over the diet periods as an indicator of nutrient flow to the gut and analyzed changes in microbial alpha and beta diversity. Fecal energy fell over the CR diet periods and post-diet period relative to baseline concentration as shown in **Figure 1A**. **Figure 1B** shows that alpha diversity metrics showed no significant difference in observed features, or Inverse Simpson, but evenness and Shannon showed significant decreases as early as week one of the diet period vs baseline. **Figure 1C-D** shows shifts in microbiome structure in the beta diversity metrics Bray-Curtis (P = 0.001) and Jaccard (P = 0.001). These metrics characterize different aspects of microbiome structure: Bray-Curtis is more representative of more abundant taxa while Jaccard is more representative of less abundant taxa. Both metrics showed the microbiome shifted from baseline starting in week 1 of the CR diet up to the end of the CR diet at week 6 and then returned immediately to similarity to the baseline in the post-diet period. This indicated large changes in both abundant and rare members of the gut microbiome.

**Figure 1.**
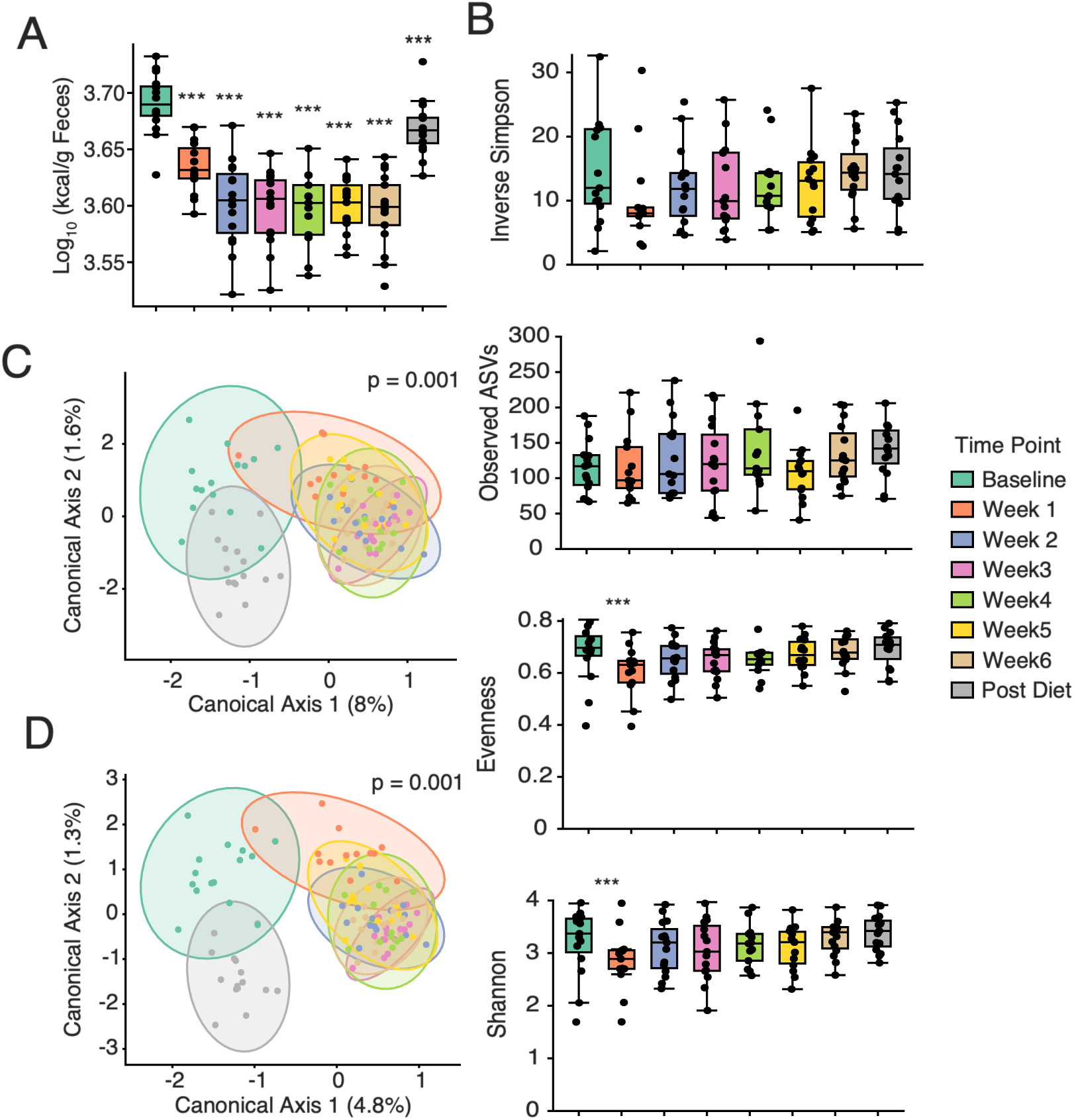
Comparison of fecal energy, alpha-diversity, and beta-diversity over the diet period relative to baseline. A) Fecal energy content was significantly lower than baseline during the CR period and the post-diet period. B)Alpha-diversity metrics. Observed genera did not change significantly. Evenness decreased significantly during week 1 of the diet relative to baseline (P = 0.041). Shannon index decreased significantly during week 1 of the diet relative to baseline (P = 0.0458). Inverse Simpson diversity did not change significantly. C) Bray-Curtis dissimilarity was significantly different (P = 0.001) from baseline at week 1 and week 6 of the diet period but returned to baseline at the post diet period. D) Jaccard similarity was also significantly different (P = 0.001) from baseline at week 1 and week 6 of the diet period but returned to baseline at the post diet period.

Communities in any ecological context display the ability to recover from perturbations in the environment^35^. This capacity is commonly referred to as “resilience”^36^. The gut microbiome is no different and displays resilience^37^. Our beta-diversity metrics support this notion. Microbiome diversity shifted significantly in weeks 1 to 6 of the CR diet likely in response to reduced nutrient flow to the colon with a swift return to baseline at the post-diet period when normal feeding resumed. These trends have been observed in other CR intervention studies^18,38,39^. However, in the CR intervention studies where participants consumed whole-foods significant changes in microbiome diversity were not observed^11,13,14,40^, suggesting CR diets may alter the microbiome differently depending on if they contain whole-foods, which reportedly feed the microbiome^10^, or not.

### Differentially abundant genera driven by reported substrate preferences and auxotrophies

We performed differential abundance using the bioinformatics tool Maaslin2 to identify genera showing significant changes in relative abundance compared to the baseline time point. Significant changes were observed in bacteria that degrade mucin or polysaccharides and bacteria with vitamin and/or amino acid auxotrophies. **Figure 2** shows the median relative abundance of taxa with an abundance of at least 1% that were differentially abundant during the CR or post-diet period compared to baseline. During the diet period, the relative abundance of ASVs representing several butyrate-producing genera decreased, including*, Gemmiger, Agathobacter, Faecalibacterium, and Agathobaculum*, taxa closely linked to butyrate production like *Bifidobacterium* also decreased. Additionally, many polysaccharide or oligosaccharide degraders such as the butyrate-producing *Monoglobus* and *Gemmiger*, as well as *Romboutsia*, *Faecalibacteria*, and *Bifidobacteria*. In contrast, the relative abundance of ASVs representing mucin degrading bacteria *Ruminococcus gnavus group* and *Mediterraneibacter*, mucin-sugar degrading bacteria *Thomasclavelia*, *and Ruthenibacterium* and the assacharolytic, amino-acid degrading *Eggerthella* increased from baseline during the CR diet period. In the post-diet period there was an increase in the relative abundance of ASVs representing bacteria with broad substrate utilization capability such as *Bacteroides* and *Achromobacter,* and rebound of butyrate producers *Faecalibacterium*, *Gemmiger*, *Agathobaculum*, and *Agathobacter*, and polysaccharide degraders *Gemmiger*, *Romboutsia* and *Monoglobus*.

**Figure 2.**
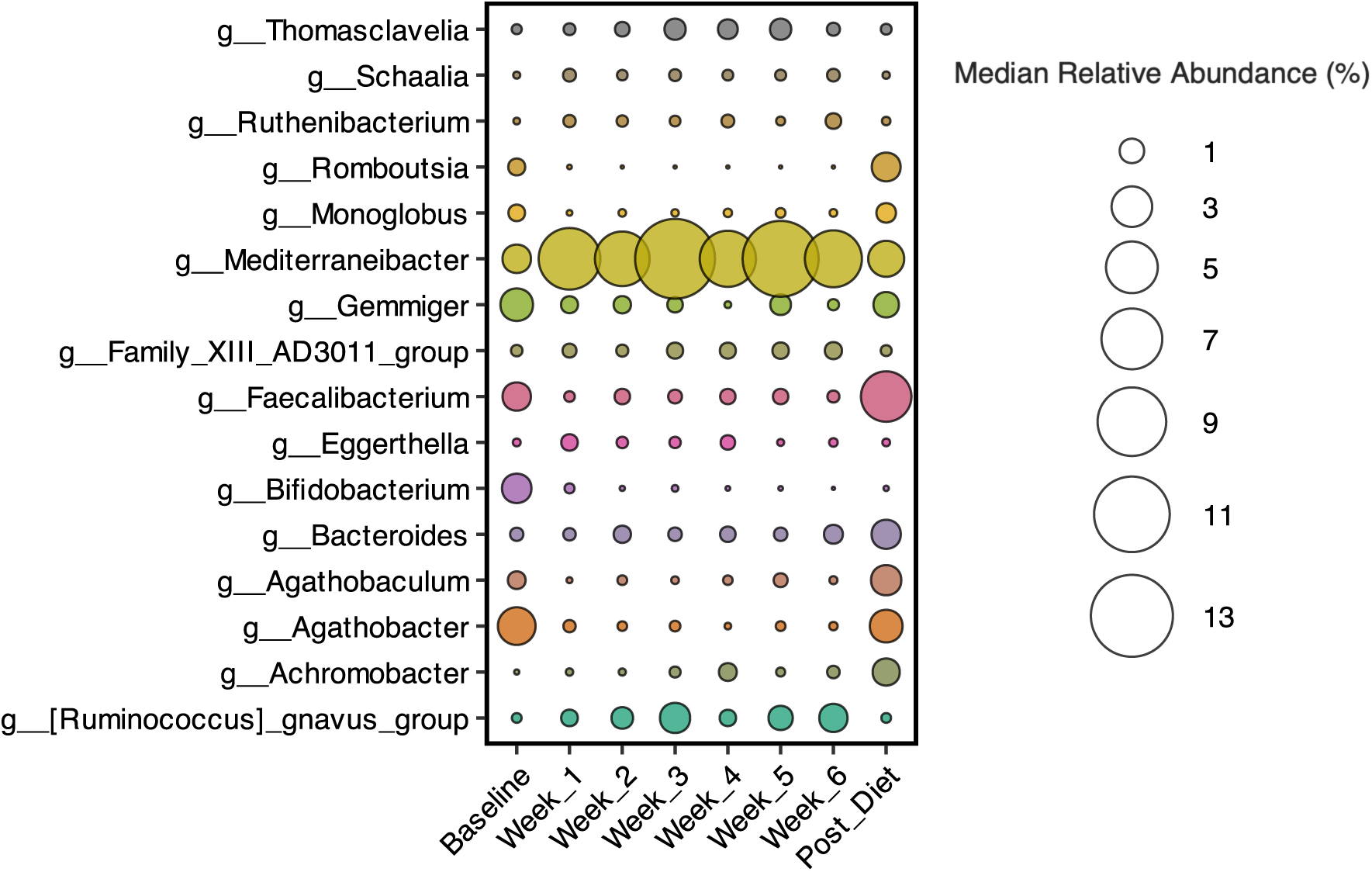
Genera with differential relative abundances compared to the baseline time point. The genera displayed had a relative abundance ≥ 1%. BH adjusted p-values were < 0.05.

The diet intervention affected the relative abundance of ASVs representing taxa closely associated with gut health. *Faecalibacterium* and *Clostridium* which decreased during the diet period, are important butyrate producers^41,42^. Butyrate producers are tightly associated with the mucin layer and important to maintaining a healthy gut epithelium and reducing inflammation^43–45^. Butyrate has been shown to increase mucus production by increasing the transcription of various host MUC genes responsible for mucin production^46,47^ as well as enzymes in the oxidative stress pathway^48^.

Many CR diets have reported an increase in *Akkermansia*, a mucin-degrading bacteria strongly associated with a healthy gut environment^21,49^. We did not see a significant change in *Akkermansia*, suggesting it was not affected by the all-liquid CR diet. However, during the diet period we saw an increase in mucin-degrading other genera *Mediterraneibacter* and *Ruminococcu*s *gnavus group* and mucin-sugar degraders *Thomasclavelia*, *and Ruthenibacterium.* This suggests a shift towards mucin utilization as an alternative energy source, as seen in other CR diets^50,51^. Unlike *Akkermansia*, *Mediterraneibacter* and *Ruminococcu*s *gnavus group* are more abundant in people who suffer from intestinal disorders such as inflammatory bowel disease (IBD)^52–54^, and irritable bowel syndrome (IBS)^55^ which are characterized by disruptions in the mucin layer and impaired gut epithelial barrier^56–58^.

Interestingly, taxa that are genetically well-equipped to degrade a range of substrates including mucins such as *Romboutsia* and *Bifidobacterium* had a lower abundance during the CR diet period compared to baseline. When looking deeper into their genomes, both bacteria are auxotrophic for either amino acids (*Bifidobacterium*) or amino acids and vitamins (*Rombutsia*). This contrasts with *Mediterraneibacter* and *Ruminococcu*s *gnavus group* which lack these auxotrophies and can even serve as providers. However, the increased relative abundance of *Eggerthella*, which has vitamin auxotrophies and a very restricted metabolism in terms of substrates, during the CR diet period raises the question of how inter-species relationships help bacteria adapt to changing environmental conditions like substrate availability.

### Predictive function analysis showed a shift in carbohydrate and amino acid metabolism and energy production during the CR diet period

Given the stratification of genera abundance on substrate preference and auxotrophies, we performed a predicted functional analysis to understand other metabolic changes that may have occurred under the CR diet. First, we used PICRUST2 to produce predicted pathways and then performed differential abundance testing with Maaslin2 to find what pathways changed during the CR diet period. Figure 3 shows pathways that were differentially abundance on 3 or more weeks during the CR diet period. The first group of pathways were more abundant during the CR diet period compared to baseline. This group includes amino acid biosynthesis pathways for arginine, sulfur amino acids, and alanine, degradation pathways for the sugars arabinose, rhamnose, fucose, the carboxylic acid ascorbate, and purine ribonucleotides, biosynthesis for the cofactors tetrahydrofolate and pyridoxyl 5’-phosphate, and biosynthesis for a glycolysis pathway. The second group shows pathways with decreased abundance during the diet period compared to baseline. These pathways include biosynthesis of polyamines, the cofactors NAD, flavin, coenzyme A, and pyrimidine deoxyribonucleotides, and degradation of the carbohydrate beta-(1,4)-mannan.

**Figure 3.**
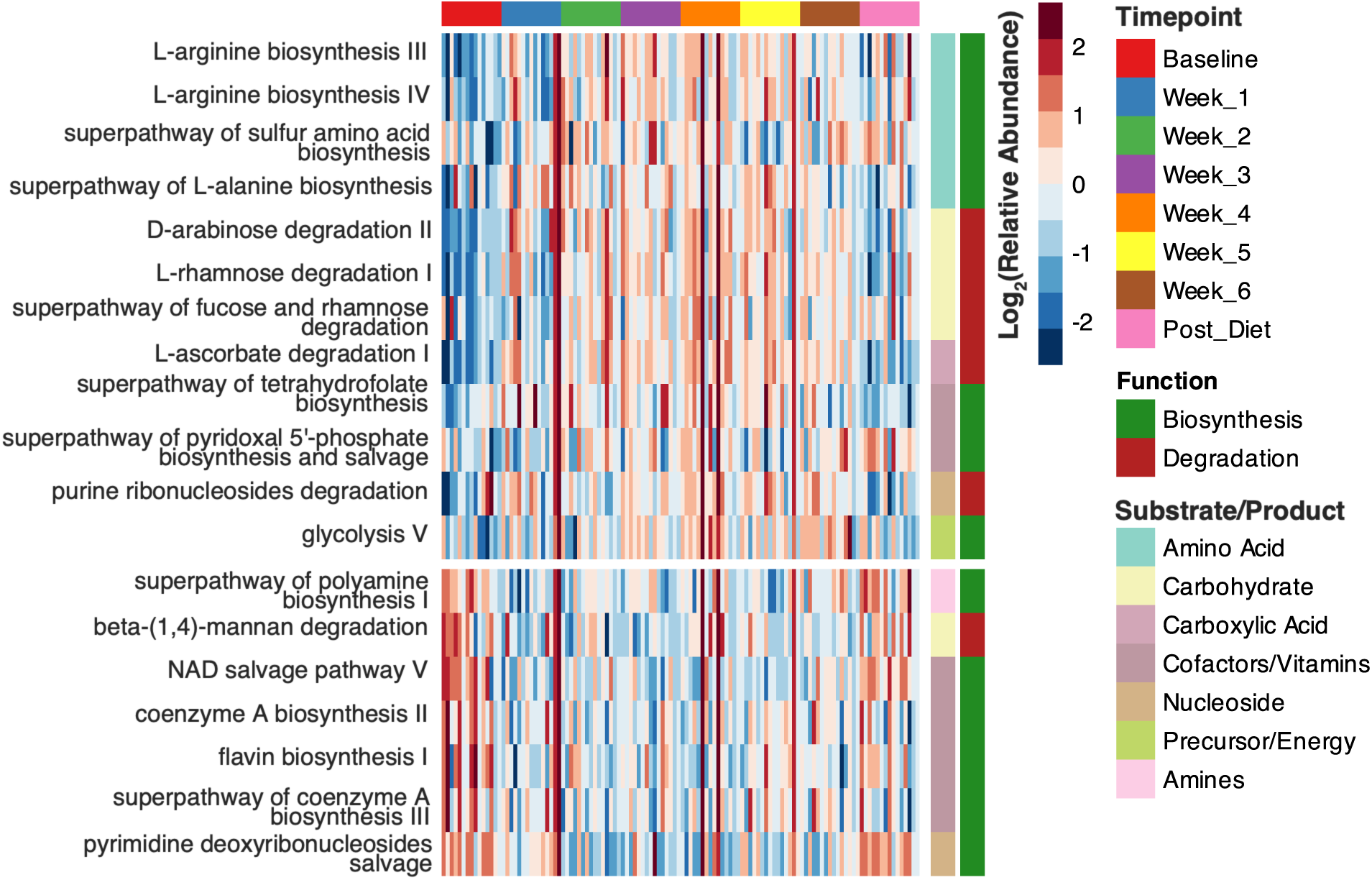
Summary of predicted MetaCyc pathways that were differentially abundant compared to baseline. MetaCyc pathways that were differentially abundant compared to baseline in at least 3 days during the diet period or only at the post diet period. All significant periods had an unadjusted p-value < 0.05 and an adjusted p-value < 0.05.

The pattern in pathway abundance suggests a shift in substrate degradation and energy metabolism. Arabinose, rhamnose, and fucose are alternative sugar substrates when glucose is scarce. Fucose is an abundant component of host-glycans and is necessary for activation of the fucose and rhamnose degradation superpathway. Another carbon source that bacteria utilize when glucose is scares is purines. Beta-(1,4)-mannan is a component of hemicellulose in plants and the decrease in degradation pathways for this polysaccharide reflect decreased abundance of polysaccharide-degrading bacterial genera. Degradation pathways aren’t the only pathways that bacteria alter during starvation conditions. They also alter biosynthesis of cofactors important for energy generation such as NAD and coenzyme A.

### Abundance of predicted genes for carbohydrate-degrading enzymes shifted based on substrate availability

The relative abundance of carbohydrate-degrading bacteria differed during the CR diet period depending on their reported substrate; host-glycan-degrading bacteria increased while polysaccharide-degrading bacteria decreased. This prompted us to assess the abundance of predicted carbohydrate-degrading genes. **Figure 4** shows the predicted mucin– and polysaccharide-degrading genes that were differentially abundant compared to baseline during the CR diet period. In **Figure 4A**, predicted genes for host-glycan-degrading α-L-fucosidase, pectin-degrading α-L-rhamnosidase, starch-degrading isoamylase and glucan 1,6-α-glucosidase, were elevated during the CR diet period. **Figure 4B** show many predicted genes for xylan, and pectin dropped during the CR diet period. Although α-L-fucosidase was the only predicted gene for a mucin-degrading enzyme differentially abundant during the diet period, predicted genes for other mucin-degrading enzymes such as protein o-glucloseaminidase, exo-alpha-sialidase, and endo-alpha-N-acetylgalactosaminidase were not differentially abundant from baseline during the CR diet (Supplemental Data 1).

**Figure 4.**
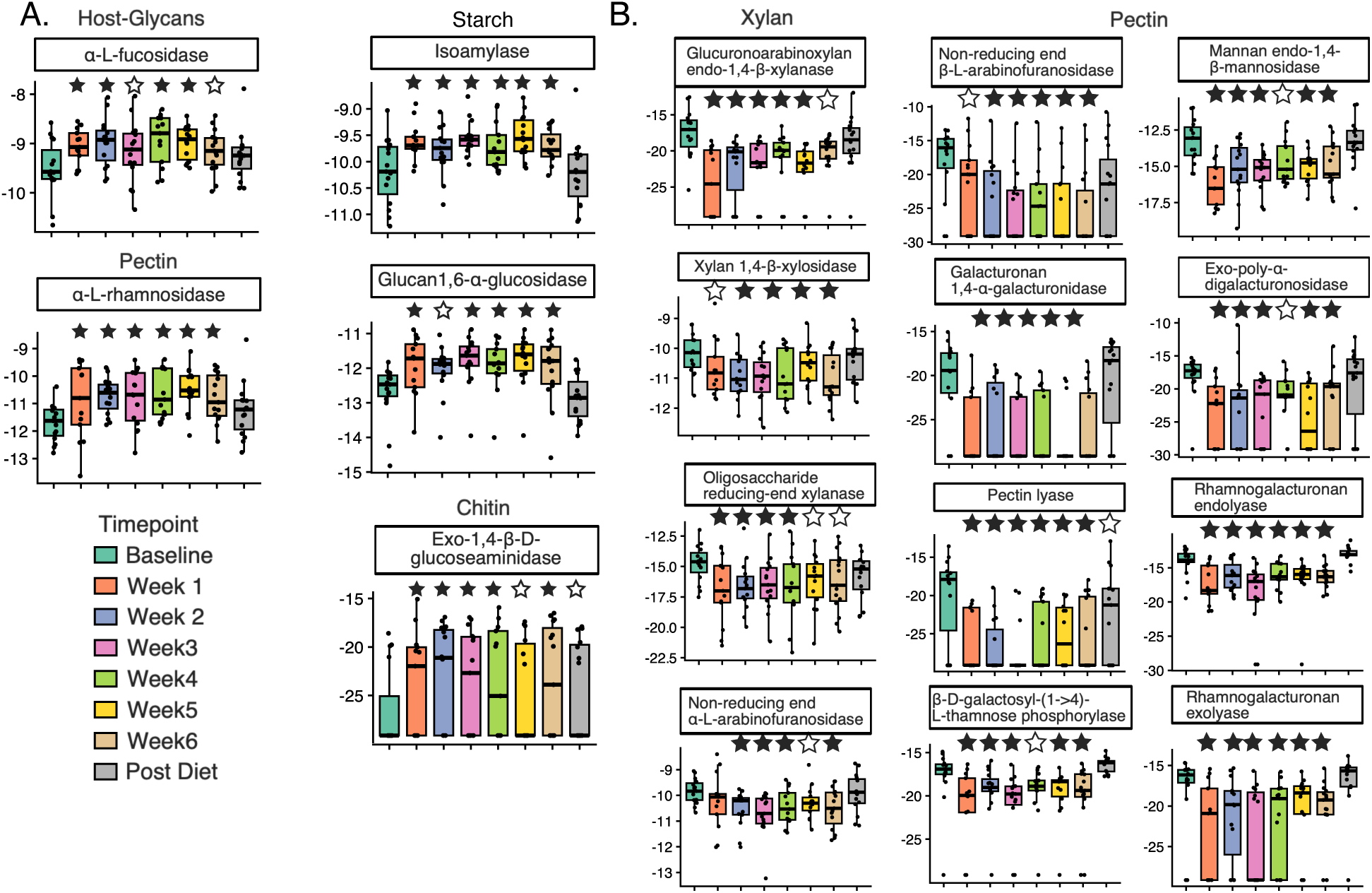
Predicted carbohydrate-degrading genes that were differentially abundant during the CR diet period compared to baseline. A) Carbohydrate-degrading genes that were higher than baseline during the CR diet period. B) Carbohydrate-degrading genes that were higher than baseline during the CR diet period. Timepoints marked with an empty star had an unadjusted p-value < 0.05. Timepoints marked with a solid star had an adjusted p-value < 0.05. Timepoints without a star had an unadjusted and adjusted p-value > 0.05.

Higher abundance of reported mucin-degrading bacteria, and lower abundance of polysaccharide-degrading genes with unchanged abundance of mucin-degrading genes indicates mucin degradation as an alternative metabolic strategy during the diet.. A metabolic shift towards host-derived glycans is also seen in other CR diet-based study^59^. This shift towards host-derived glycans is likely in response to the low carbohydrate availability in the colon with a recovery of diet-derived carbohydrate preference likely due to a return to a diet higher in polysaccharides by the post-diet period.

### Predicted genes involved in butyrate-production suggest a shift in substrate from carbohydrates to the amino acid lysine

Bacteria can ferment different substrates, such as sugars, carboxylates, and amino acids, to produce butyrate. We performed differential abundance testing on predicted genes involved in butyrate production to decipher changes in butyrate producing metabolism. **Figure 5** shows the predicted genes that showed differential abundance during the CR period compared to the baseline. **Figure 5A** shows the predicted genes that were elevated during the CR period compared to baseline. The first two genes, L-erythro-3,5-diaminohexanoate dehydrogenase and Lysine 5,6-aminomutase, are involved in the pathway that ferments the amino acid L-lysine to produce butyrate. Acetyl-CoA C-acetyltransferase converts acetyl-CoA into acetoacetyl-CoA which is then converted to butyrate through reverse β-oxidation. Lastly, acetate—CoA ligase (ADP-forming) redirects acetate into acetyl-CoA which can then be utilized produce butyrate. **Figure 5B** shows the two predicted genes, phosphate butyryltransferase and butyrate kinase, that were reduced during the CR period compared to the baseline. The enzyme products of these genes convert butyryl-CoA into butyrate.

**Figure 5.**
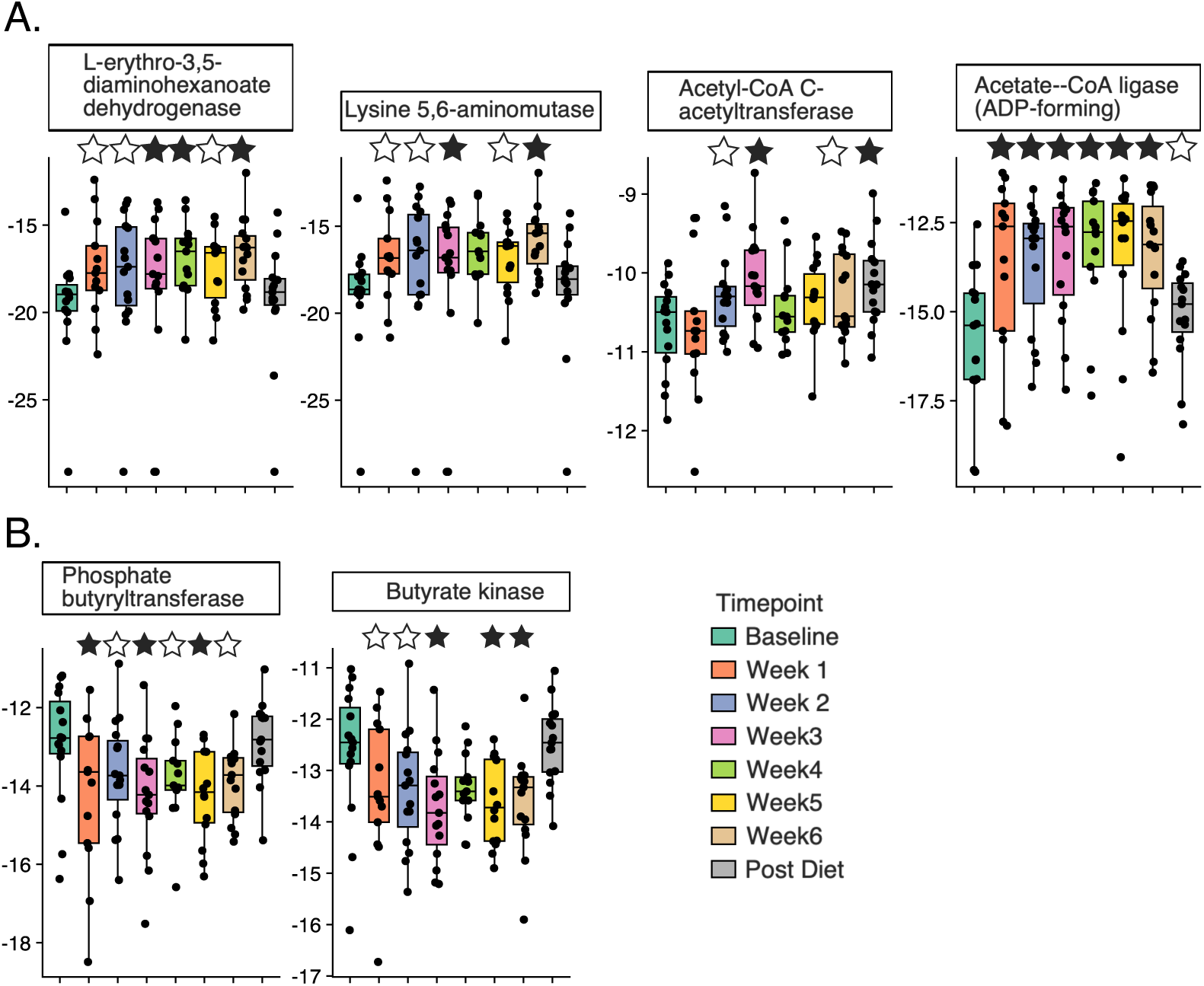
Differentially abundant predicted genes involved in butyrate production. A) Predicted genes involved in butyrate production that were higher during the CR period compared to baseline. B) Predicted genes involved in butyrate production that were lower during the CR period compared to baseline. Timepoints marked with an empty star had an unadjusted p-value < 0.05. Timepoints marked with a solid star had an adjusted p-value < 0.05. Timepoints without a star had an unadjusted and adjusted p-value > 0.05.

These changes in predicted gene function suggest that gut microbial community functions changed by changes in abundant bacteria, for example butyrate-producing bacteria with different metabolic pathways shifted during the CR diet. One shift in the presence of bacteria, is an increase in bacteria that ferment amino acids, specifically L-lysine to produce butyrate. This change in community metabolism is likely an adaptation to the reduced overall availability of dietary carbohydrates in the all-liquid CR diet. Carbohydrates are the preferred source for energy and carbon for gut microbiota. Once dietary carbohydrates are consumed, the microbiota then start to degrade amino acids for energy and carbon^60^, and microbiota good at this function, will be enriched. The other shift was in the terminal pathway of butyrate production. Once crotonoyl-CoA is converted to butyryl-CoA, butyrate can be produced in one of two ways. The first is through CoA transferases that transfer the CoA to acetoacetate or acetate yielding butyrate. This is the most common pathway found in the human gut microbiome^61^. The second pathway replaces the CoA with a phosphate then a kinase transfers the phosphate to an ADP yielding ATP and butyrate. This pathway is less common in the human gut microbiome^61^. Some butyrate-producing genera that had a lower abundance during the CR diet period utilize this pathway, such as *Eubacterium ventrosum* group, *Clostridium*, and *Gemmiger*^62^. We hypothesize that the carbohydrates in the all-liquid CR diet were absorbed by the host leaving little, if any, carbohydrates entering the colon to feed the microbiota. And although there was a decrease in butyrate producers, especially those that feed on polysaccharides or possess multiple auxotrophies, other butyrate producers that feed on alternative carbon-sources, such as mucin like *Thomasclavelia*, were able to thrive.

## Conclusion

This analysis demonstrates that an all-liquid CR diet induced significant shifts in the gut microbiome community and predicted function. While alpha-diversity did not show any significant changes beyond the first week of diet intervention, beta-diversity showed a consistent shift in both high and low abundance bacteria. The microbiome community composition showed alterations specifically among butyrate-producing and mucin-degrading bacteria While butyrate-producing bacteria declined, mucin degraders increased, suggesting a shift in microbial metabolism during and after the diet. Predicted functional pathway analysis revealed a temporary reduction (within diet period) in gene abundance for carbohydrate fermentation and an increase for amino acid metabolism reflecting microbial adaptation to nutrient availability. These findings underscore the resilience of the gut microbiome but also highlight potentially detrimental health implications, particularly regarding mucin degradation and gut permeability.

Our results also highlight some important differences between an all-liquid CR diet and other CR diets that incorporate whole foods. One difference was that CR diets with whole foods generally had less of an effect on microbial beta diversity than our study. Another difference was the increased abundance of mucin-degrading bacteria strongly associated with impaired gut health. Other CR diets observe an increase in mucin degraders, but they tend to be bacteria strongly associated with a healthy gut. These important differences suggest that some of the changes we reported are common to CR diets with whole food components, but all-liquid CR diets had a more significant impact on bacterial community structure and encouraged the growth of bacteria potentially detrimental to gut health. Although all-liquid CR diets are used for a variety of health-related purposes, supplementation of these diets with dietary fiber, when appropriate, may reduce the risk of harmful health effects by encouraging the growth of beneficial bacteria, though more research is needed.

A weakness of this study is that it is a longitudinal study of just one diet intervention: the all-liquid CR diet. Future studies could include a whole-food CR diet on a group of similar people and compare how CR diet composition affects the gut microbiota and the host in general and in terms of gut barrier health and function. These studies could also explore the effects of a CR diet on different populations to better understand how interindividual variation influences host response to a CR diet and how the gut microbiome contributes to that host response. This study also had a small sample size (N = 12). Future studies with a larger sample size are needed to confirm our findings here. Another weakness of this study is that it only included data from 16S rRNA gene amplicon sequencing which limited community analysis to the genera level and function analysis to functional potential from predictions based on genomes. Using shot gun metagenomic sequencing would provide greater resolution to the species level and RNA sequencing would provide a more accurate picture of the microbial metabolism response to CR diet interventions.

## Author Contributions

S.V. and J.K. designed the study. B.D. performed the sequencing analysis, statistical analysis, and wrote the manuscript. K.N., R.K.B, S.V., T.H., and J.K. provided critical revision and input during drafting of the manuscript.

## Availability of Data

The data presented in this study are openly available in the NCBI SRA repository under BioProject ID PRJNA1565460 and can be accessed at https://www.ncbi.nlm.nih.gov/bioproject/1565460.

## Supplementary Material

**Supplementary Table 1.**
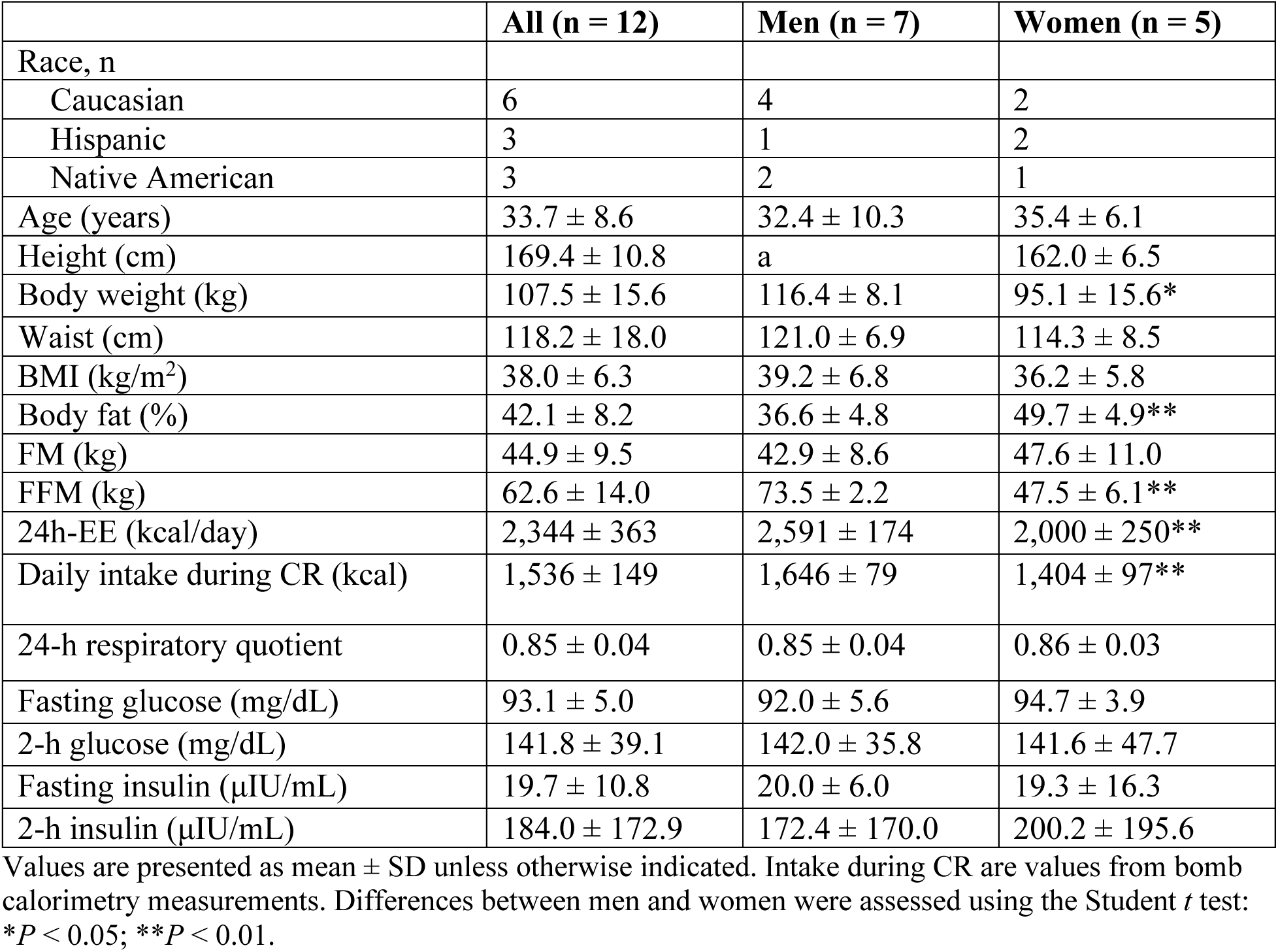
Table 1 General characteristics of the study participants [1].

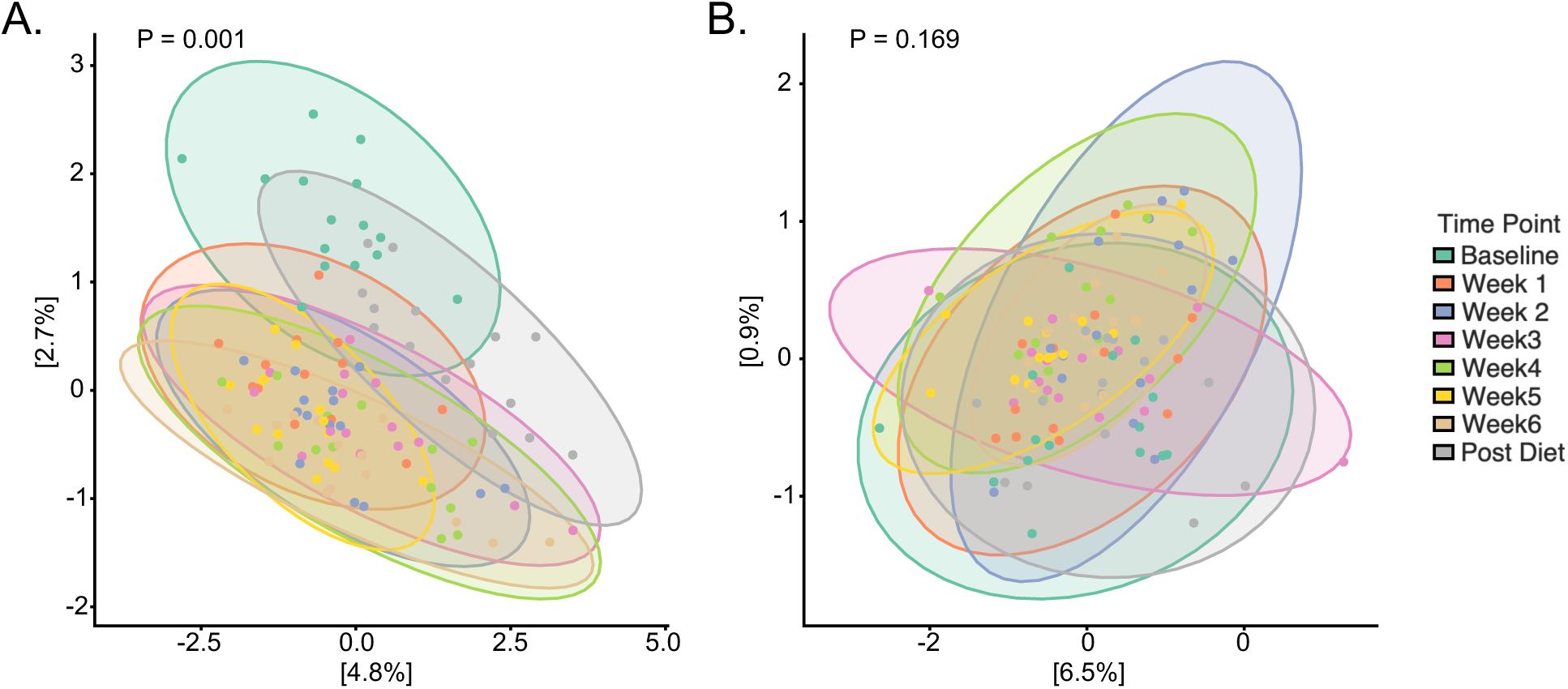
Supplementary Figure 1.

**Supplementary Figure 2.** Unweighted and weighted Unifrac metrics for beta-diversity. A) Unweighted unifrac was different by diet period (Permanova test, P = 0.001). B) Weighted unifrac was not different by diet period (Permanova test, P = 0.169).

---

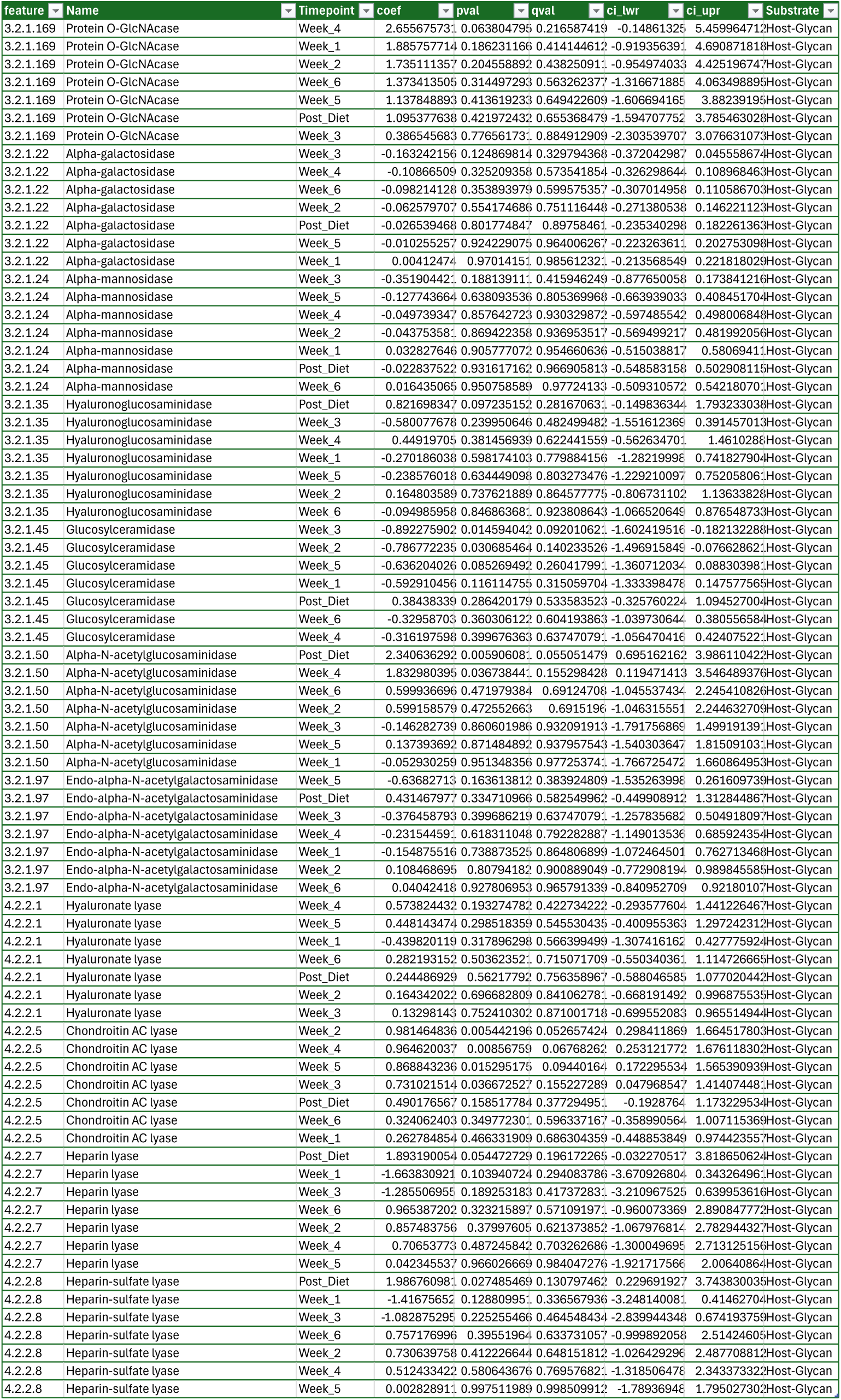
Supplementary Data 1.

| feature | Name | Timepoint | coef | pval | qval | ci_lwr | ci_upr | Substrate |
|---|---|---|---|---|---|---|---|---|
| 3.2.1.169 | Protein O-GlcNAcase | Week_4 | 2.655675731 | 0.063804795 | 0.216587419 | -0.14861325 | 5.459964712 | Host-Glycan |
| 3.2.1.169 | Protein O-GlcNAcase | Week_1 | 1.885757714 | 0.186231166 | 0.414144612 | -0.919356391 | 4.690871818 | Host-Glycan |
| 3.2.1.169 | Protein O-GlcNAcase | Week_2 | 1.735111357 | 0.204558892 | 0.438250911 | -0.954974033 | 4.425196747 | Host-Glycan |
| 3.2.1.169 | Protein O-GlcNAcase | Week_6 | 1.373413505 | 0.314497293 | 0.563262377 | -1.316671885 | 4.063498895 | Host-Glycan |
| 3.2.1.169 | Protein O-GlcNAcase | Week_5 | 1.137848893 | 0.413619233 | 0.649422609 | -1.606694165 | 3.88239195 | Host-Glycan |
| 3.2.1.169 | Protein O-GlcNAcase | Post_Diet | 1.095377638 | 0.421972432 | 0.655368479 | -1.594707752 | 3.785463028 | Host-Glycan |
| 3.2.1.169 | Protein O-GlcNAcase | Week_3 | 0.386545683 | 0.776561731 | 0.884912909 | -2.303539707 | 3.076631073 | Host-Glycan |
| 3.2.1.22 | Alpha-galactosidase | Week_3 | -0.163242156 | 0.124869814 | 0.329794368 | -0.372042987 | 0.045558674 | Host-Glycan |
| 3.2.1.22 | Alpha-galactosidase | Week_4 | -0.10866509 | 0.325209358 | 0.573541854 | -0.326298644 | 0.108968463 | Host-Glycan |
| 3.2.1.22 | Alpha-galactosidase | Week_6 | -0.098214128 | 0.353893979 | 0.599575357 | -0.307014958 | 0.110586703 | Host-Glycan |
| 3.2.1.22 | Alpha-galactosidase | Week_2 | -0.062579707 | 0.554174686 | 0.751116448 | -0.271380538 | 0.146221123 | Host-Glycan |
| 3.2.1.22 | Alpha-galactosidase | Post_Diet | -0.026539468 | 0.801774847 | 0.89758461 | -0.235340298 | 0.182261363 | Host-Glycan |
| 3.2.1.22 | Alpha-galactosidase | Week_5 | -0.010255257 | 0.924229075 | 0.964006267 | -0.223263611 | 0.202753098 | Host-Glycan |
| 3.2.1.22 | Alpha-galactosidase | Week_1 | 0.00412474 | 0.97014151 | 0.985612321 | -0.213568549 | 0.221818029 | Host-Glycan |
| 3.2.1.24 | Alpha-mannosidase | Week_3 | -0.351904421 | 0.188139111 | 0.415946249 | -0.877650058 | 0.173841216 | Host-Glycan |
| 3.2.1.24 | Alpha-mannosidase | Week_5 | -0.127743664 | 0.638093536 | 0.805369968 | -0.663939033 | 0.408451704 | Host-Glycan |
| 3.2.1.24 | Alpha-mannosidase | Week_4 | -0.049739347 | 0.857642723 | 0.930329872 | -0.597485542 | 0.498006848 | Host-Glycan |
| 3.2.1.24 | Alpha-mannosidase | Week_2 | -0.043753581 | 0.869422358 | 0.936953517 | -0.569499217 | 0.481992056 | Host-Glycan |
| 3.2.1.24 | Alpha-mannosidase | Week_1 | 0.032827646 | 0.905777072 | 0.954660636 | -0.515038817 | 0.58069411 | Host-Glycan |
| 3.2.1.24 | Alpha-mannosidase | Post_Diet | -0.022837522 | 0.931617162 | 0.966905813 | -0.548583158 | 0.502908115 | Host-Glycan |
| 3.2.1.24 | Alpha-mannosidase | Week_6 | 0.016435065 | 0.950758589 | 0.97724133 | -0.509310572 | 0.542180701 | Host-Glycan |
| 3.2.1.35 | Hyaluronoglucosaminidase | Post_Diet | 0.821698347 | 0.097235152 | 0.28167063 | -0.149836344 | 1.793233038 | Host-Glycan |
| 3.2.1.35 | Hyaluronoglucosaminidase | Week_3 | -0.580077678 | 0.239950646 | 0.482499482 | -1.551612369 | 0.391457013 | Host-Glycan |
| 3.2.1.35 | Hyaluronoglucosaminidase | Week_4 | 0.44919705 | 0.381456939 | 0.622441559 | -0.562634701 | 1.4610288 | Host-Glycan |
| 3.2.1.35 | Hyaluronoglucosaminidase | Week_1 | -0.270186038 | 0.598174103 | 0.779884156 | -1.28219998 | 0.741827904 | Host-Glycan |
| 3.2.1.35 | Hyaluronoglucosaminidase | Week_5 | -0.238576018 | 0.634449098 | 0.803273476 | -1.229210097 | 0.752058061 | Host-Glycan |
| 3.2.1.35 | Hyaluronoglucosaminidase | Week_2 | 0.164803589 | 0.737621889 | 0.864577775 | -0.806731102 | 1.13633828 | Host-Glycan |
| 3.2.1.35 | Hyaluronoglucosaminidase | Week_6 | -0.094985958 | 0.846863681 | 0.923808643 | -1.066520649 | 0.876548733 | Host-Glycan |
| 3.2.1.45 | Glucosylceramidase | Week_3 | -0.892275902 | 0.014594042 | 0.092010621 | -1.602419516 | -0.18213228 | Host-Glycan |
| 3.2.1.45 | Glucosylceramidase | Week_2 | -0.786772235 | 0.030685464 | 0.140233526 | -1.496915849 | -0.076628621 | Host-Glycan |
| 3.2.1.45 | Glucosylceramidase | Week_5 | -0.636204026 | 0.085269492 | 0.260417991 | -1.360712034 | 0.088303981 | Host-Glycan |
| 3.2.1.45 | Glucosylceramidase | Week_1 | -0.592910456 | 0.116114755 | 0.315059704 | -1.333398478 | 0.147577565 | Host-Glycan |
| 3.2.1.45 | Glucosylceramidase | Post_Diet | 0.38438339 | 0.286420179 | 0.533583523 | -0.325760224 | 1.094527004 | Host-Glycan |
| 3.2.1.45 | Glucosylceramidase | Week_6 | -0.32958703 | 0.360306122 | 0.604193863 | -1.039730644 | 0.380556584 | Host-Glycan |
| 3.2.1.45 | Glucosylceramidase | Week_4 | -0.316197598 | 0.399676363 | 0.637470791 | -1.056470416 | 0.424075221 | Host-Glycan |
| 3.2.1.50 | Alpha-N-acetylglucosaminidase | Post_Diet | 2.340636292 | 0.005906081 | 0.055051479 | 0.695162162 | 3.986110422 | Host-Glycan |
| 3.2.1.50 | Alpha-N-acetylglucosaminidase | Week_4 | 1.832980395 | 0.036738441 | 0.155298428 | 0.119471413 | 3.546489376 | Host-Glycan |
| 3.2.1.50 | Alpha-N-acetylglucosaminidase | Week_6 | 0.599936696 | 0.471979384 | 0.69124708 | -1.045537434 | 2.245410826 | Host-Glycan |
| 3.2.1.50 | Alpha-N-acetylglucosaminidase | Week_2 | 0.599158579 | 0.472552663 | 0.6915196 | -1.046315551 | 2.244632709 | Host-Glycan |
| 3.2.1.50 | Alpha-N-acetylglucosaminidase | Week_3 | -0.146282739 | 0.860601986 | 0.932091913 | -1.791756869 | 1.499191391 | Host-Glycan |
| 3.2.1.50 | Alpha-N-acetylglucosaminidase | Week_5 | 0.137393692 | 0.871484892 | 0.937957543 | -1.540303647 | 1.815091031 | Host-Glycan |
| 3.2.1.50 | Alpha-N-acetylglucosaminidase | Week_1 | -0.052930259 | 0.951348356 | 0.977253741 | -1.766725472 | 1.660864953 | Host-Glycan |
| 3.2.1.97 | Endo-alpha-N-acetylgalactosaminidase | Week_5 | -0.63682713 | 0.163613812 | 0.383924809 | -1.535263998 | 0.261609739 | Host-Glycan |
| 3.2.1.97 | Endo-alpha-N-acetylgalactosaminidase | Post_Diet | 0.431467977 | 0.334710966 | 0.582549962 | -0.449908912 | 1.312844867 | Host-Glycan |
| 3.2.1.97 | Endo-alpha-N-acetylgalactosaminidase | Week_3 | -0.376458793 | 0.399686219 | 0.637470791 | -1.257835682 | 0.504918097 | Host-Glycan |
| 3.2.1.97 | Endo-alpha-N-acetylgalactosaminidase | Week_4 | -0.231544591 | 0.618311048 | 0.792282887 | -1.149013536 | 0.685924354 | Host-Glycan |
| 3.2.1.97 | Endo-alpha-N-acetylgalactosaminidase | Week_1 | -0.154875516 | 0.738873525 | 0.864806899 | -1.072464501 | 0.762713468 | Host-Glycan |
| 3.2.1.97 | Endo-alpha-N-acetylgalactosaminidase | Week_2 | 0.108468695 | 0.80794182 | 0.900889049 | -0.772908194 | 0.989845585 | Host-Glycan |
| 3.2.1.97 | Endo-alpha-N-acetylgalactosaminidase | Week_6 | 0.04042418 | 0.927806953 | 0.965791339 | -0.840952709 | 0.92180107 | Host-Glycan |
| 4.2.2.1 | Hyaluronate lyase | Week_4 | 0.573824432 | 0.193274782 | 0.422734222 | -0.293577604 | 1.441226467 | Host-Glycan |
| 4.2.2.1 | Hyaluronate lyase | Week_5 | 0.448143474 | 0.298518359 | 0.545530435 | -0.400955363 | 1.297242312 | Host-Glycan |
| 4.2.2.1 | Hyaluronate lyase | Week_1 | -0.439820119 | 0.317896298 | 0.566399499 | -1.307416162 | 0.427775924 | Host-Glycan |
| 4.2.2.1 | Hyaluronate lyase | Week_6 | 0.282193152 | 0.503623521 | 0.715071709 | -0.550340361 | 1.114726665 | Host-Glycan |
| 4.2.2.1 | Hyaluronate lyase | Post_Diet | 0.244486929 | 0.56217792 | 0.756358967 | -0.588046585 | 1.077020442 | Host-Glycan |
| 4.2.2.1 | Hyaluronate lyase | Week_2 | 0.164342022 | 0.696682809 | 0.841062781 | -0.668191492 | 0.996875535 | Host-Glycan |
| 4.2.2.1 | Hyaluronate lyase | Week_3 | 0.13298143 | 0.752410302 | 0.871001718 | -0.699552083 | 0.965514944 | Host-Glycan |
| 4.2.2.5 | Chondroitin AC lyase | Week_2 | 0.981464836 | 0.005442196 | 0.052657424 | 0.298411869 | 1.664517803 | Host-Glycan |
| 4.2.2.5 | Chondroitin AC lyase | Week_4 | 0.964620037 | 0.00856759 | 0.06768262 | 0.253121772 | 1.676118302 | Host-Glycan |
| 4.2.2.5 | Chondroitin AC lyase | Week_5 | 0.868843236 | 0.015295175 | 0.09440164 | 0.172295534 | 1.565390939 | Host-Glycan |
| 4.2.2.5 | Chondroitin AC lyase | Week_3 | 0.731021514 | 0.036672527 | 0.155227289 | 0.047968547 | 1.414074481 | Host-Glycan |
| 4.2.2.5 | Chondroitin AC lyase | Post_Diet | 0.490176567 | 0.158517784 | 0.377294951 | -0.1928764 | 1.173229534 | Host-Glycan |
| 4.2.2.5 | Chondroitin AC lyase | Week_6 | 0.324062403 | 0.349772301 | 0.596337167 | -0.358990564 | 1.007115369 | Host-Glycan |
| 4.2.2.5 | Chondroitin AC lyase | Week_1 | 0.262784854 | 0.466331909 | 0.686304359 | -0.448853849 | 0.974423557 | Host-Glycan |
| 4.2.2.7 | Heparin lyase | Post_Diet | 1.893190054 | 0.054472729 | 0.196172265 | -0.032270517 | 3.818650624 | Host-Glycan |
| 4.2.2.7 | Heparin lyase | Week_1 | -1.663830921 | 0.103940724 | 0.294083786 | -3.670926804 | 0.343264961 | Host-Glycan |
| 4.2.2.7 | Heparin lyase | Week_3 | -1.285506955 | 0.189253183 | 0.417372831 | -3.210967525 | 0.639953616 | Host-Glycan |
| 4.2.2.7 | Heparin lyase | Week_6 | 0.965387202 | 0.323215897 | 0.571091971 | -0.960073369 | 2.890847772 | Host-Glycan |
| 4.2.2.7 | Heparin lyase | Week_2 | 0.857483756 | 0.37997605 | 0.621373852 | -1.067976814 | 2.782944327 | Host-Glycan |
| 4.2.2.7 | Heparin lyase | Week_4 | 0.70653773 | 0.487245842 | 0.703262686 | -1.300049695 | 2.713125156 | Host-Glycan |
| 4.2.2.7 | Heparin lyase | Week_5 | 0.042345537 | 0.966026669 | 0.984047276 | -1.921717566 | 2.00640864 | Host-Glycan |
| 4.2.2.8 | Heparin-sulfate lyase | Post_Diet | 1.986760981 | 0.027485469 | 0.130797462 | 0.229691927 | 3.743830035 | Host-Glycan |
| 4.2.2.8 | Heparin-sulfate lyase | Week_1 | -1.41675652 | 0.128809951 | 0.336567936 | -3.248140081 | 0.41462704 | Host-Glycan |
| 4.2.2.8 | Heparin-sulfate lyase | Week_3 | -1.082875295 | 0.225255466 | 0.464548434 | -2.839944348 | 0.674193759 | Host-Glycan |
| 4.2.2.8 | Heparin-sulfate lyase | Week_6 | 0.757176996 | 0.39551964 | 0.633731057 | -0.999892058 | 2.51424605 | Host-Glycan |
| 4.2.2.8 | Heparin-sulfate lyase | Week_2 | 0.730639758 | 0.412226644 | 0.648151812 | -1.026429296 | 2.487708812 | Host-Glycan |
| 4.2.2.8 | Heparin-sulfate lyase | Week_4 | 0.512433422 | 0.580643676 | 0.769576821 | -1.318506478 | 2.343373322 | Host-Glycan |
| 4.2.2.8 | Heparin-sulfate lyase | Week_5 | 0.002828911 | 0.997511989 | 0.998509912 | -1.78936948 | 1.795027302 | Host-Glycan |

## References

1. Tanca, A. et al. Caloric restriction promotes functional changes involving short-chain fatty acid biosynthesis in the rat gut microbiota. Sci. Rep. 8, 1–13 (2018).

2. Bartley, J. M., Zhou, X., Kuchel, G. A., Weinstock, G. M. & Haynes, L. Impact of Age, Caloric Restriction, and Influenza Infection on Mouse Gut Microbiome: An Exploratory Study of the Role of Age-Related Microbiome Changes on Influenza Responses. Front. Immunol. 8, (2017).

3. Benítez-Páez, A. et al. Sex, Food, and the Gut Microbiota: Disparate Response to Caloric Restriction Diet with Fiber Supplementation in Women and Men. Mol. Nutr. Food Res. 65, 1–13 (2021).

4. Anton, S. & Leeuwenburgh, C. Fasting or caloric restriction for healthy aging. Exp. Gerontol. 48, 1003–1005 (2013).

5. Masoro, E. J. Overview of caloric restriction and ageing. Mech. Ageing Dev. 126, 913–922 (2005).

6. Kebbe, M., Sparks, J. R., Flanagan, E. W. & Redman, L. M. Beyond weight loss: current perspectives on the impact of calorie restriction on healthspan and lifespan. Expert Rev. Endocrinol. Metab. 16, 95–108 (2021).

7. Spadaro, O. et al. Caloric restriction in humans reveals immunometabolic regulators of health span. Science (1979). 375, 671–677 (2022).

8. Mohr, A. E. et al. Gut microbiome remodeling and metabolomic profile improves in response to protein pacing with intermittent fasting versus continuous caloric restriction. Nat. Commun. 15, 1– 20 (2024).

9. Asnicar, F. et al. Microbiome connections with host metabolism and habitual diet from 1,098 deeply phenotyped individuals. Nat. Med. 27, 321–332 (2021).

10. Corbin, K. D. et al. Host-diet-gut microbiome interactions influence human energy balance: a randomized clinical trial. Nat. Commun. 14, 3161 (2023).

11. Ott, B. et al. Effect of caloric restriction on gut permeability, inflammation markers, and fecal microbiota in obese women. Sci. Rep. 7, 1–10 (2017).

12. Zou, H. et al. Effect of Caloric Restriction on BMI, Gut Microbiota, and Blood Amino Acid Levels in Non-Obese Adults. Nutrients 12, 631 (2020).

13. Mohr, A. E. et al. Gut microbiome remodeling and metabolomic profile improves in response to protein pacing with intermittent fasting versus continuous caloric restriction. Nat. Commun. 15, 1– 20 (2024).

14. Dong, T. S. et al. A high protein calorie restriction diet alters the gut microbiome in obesity. Nutrients 12, 1–16 (2020).

15. Maifeld, A. et al. Fasting alters the gut microbiome reducing blood pressure and body weight in metabolic syndrome patients. Nat. Commun. 12, (2021).

16. Dao, M. C. et al. Akkermansia muciniphila and improved metabolic health during a dietary intervention in obesity: Relationship with gut microbiome richness and ecology. Gut 65, 426–436 (2016).

17. Kern, L., Kviatcovsky, D., He, Y. & Elinav, E. Impact of caloric restriction on the gut microbiota. Curr. Opin. Microbiol. 73, 1–12 (2023).

18. von Schwartzenberg, R. J. et al. Caloric restriction disrupts the microbiota and colonization resistance. Nature 595, 272–277 (2021).

19. Murray, D. P., Welsh, J. D., Rankin, R. A. & Warner, R. Survey: Use of clear and full liquid diets with or without commercially produced formulas. Journal of Parenteral and Enteral Nutrition 9, 732–734 (1985).

20. Chao, A. M., Quigley, K. M. & Wadden, T. A. Dietary interventions for obesity: Clinical and mechanistic findings. Journal of Clinical Investigation 131, (2021).

21. Lane, M. et al. The effect of ultra-processed very low-energy diets on gut microbiota and metabolic outcomes in individuals with obesity: A systematic literature review. Obes. Res. Clin. Pract. 14, 197–204 (2020).

22. Hernández-Calderón, P., Wiedemann, L. & Benítez-Páez, A. The microbiota composition drives personalized nutrition: Gut microbes as predictive biomarkers for the success of weight loss diets. Front. Nutr. 9, (2022).

23. Reinhardt, M. et al. A Human Thrifty Phenotype Associated With Less Weight Loss During Caloric Restriction. Diabetes 64, 2859–2867 (2015).

24. Caporaso, J. G. et al. Global patterns of 16S rRNA diversity at a depth of millions of sequences per sample. Proc. Natl. Acad. Sci. U. S. A. 108, 4516–4522 (2011).

25. Bolyen, E. et al. Reproducible, interactive, scalable and extensible microbiome data science using QIIME 2. Nat. Biotechnol. 37, 852–857 (2019).

26. Sorensen, T. A Method of Establishing Groups of Equal Amplitude in Plant Sociology Based on Similarity of Species Content. *Det. Kong. Danske Vidensk*, Selesk Biology Skr 5, 1–34 (1948).

27. Jaccard, P. Sur La Distribution Florale. Bulletin de la Société Vaudoise des Sciences Naturelles 44, 223–270 (1908).

28. McMurdie, P. J. & Holmes, S. Phyloseq: An R Package for Reproducible Interactive Analysis and Graphics of Microbiome Census Data. PLoS One 8, e61217 (2013).

29. Douglas, G. M. et al. PICRUSt2 for prediction of metagenome functions. Nat. Biotechnol. 38, 685–688 (2020).

30. Wickham, H. *Ggplot2: Elegant Graphics for Data Analysis*. (Springer International Publishing, Cham, 2016). doi:10.1007/978-3-319-24277-4.

31. David, L. A. et al. Diet rapidly and reproducibly alters the human gut microbiome. Nature 505, 559–563 (2014).

32. Mallick, H. et al. Multivariable association discovery in population-scale meta-omics studies. PLoS Comput. Biol. 17, e1009442 (2021).

33. Zmora, N., Suez, J. & Elinav, E. You are what you eat: diet, health and the gut microbiota. Nat. Rev. Gastroenterol. Hepatol. 16, 35–56 (2019).

34. Paine, R. T., Tegner, M. J. & Johnson, E. a. Ecological Surprises. Ecosystems 1, 535–545 (1998).

35. Walker, B., Holling, C. S., Carpenter, S. R. & Kinzig, A. Resilience, adaptability and transformability in social-ecological systems. Ecology and Society 9, (2004).

36. Fassarella, M. et al. Gut microbiome stability and resilience: Elucidating the response to perturbations in order to modulate gut health. Gut 70, 595–605 (2021).

37. Sang, X. et al. Dynamics and ecological reassembly of the human gut microbiome and the host metabolome in response to prolonged fasting. https://doi.org/10.3389/fmicb.2023.1265425 (2023) doi:10.3389/fmicb.2023.1265425.

38. Louis, S., Tappu, R. M., Damms-Machado, A., Huson, D. H. & Bischoff, S. C. Characterization of the gut microbial community of obese patients following a weight-loss intervention using whole metagenome shotgun sequencing. PLoS One 11, 1–18 (2016).

39. Sowah, S. A. et al. Calorie restriction improves metabolic state independently of gut microbiome composition: a randomized dietary intervention trial. Genome Med. 14, 1–21 (2022).

40. Tamanai-Shacoori, Z. et al. Roseburia spp.: A marker of health? Future Microbiol. 12, 157–170 (2017).

41. Miquel, S. et al. Faecalibacterium prausnitzii and human intestinal health. Curr. Opin. Microbiol. 16, 255–261 (2013).

42. Mohebali, N. et al. Faecalibacterium prausnitzii, Bacteroides faecis and Roseburia intestinalis attenuate clinical symptoms of experimental colitis by regulating Treg/Th17 cell balance and intestinal barrier integrity. Biomedicine and Pharmacotherapy 167, 115568 (2023).

43. Mohebali, N., Ekat, K., Kreikemeyer, B. & Breitrück, A. Barrier protection and recovery effects of gut commensal bacteria on differentiated intestinal epithelial cells in vitro. Nutrients 12, 1–23 (2020).

44. Singh, V. et al. Butyrate producers, “The Sentinel of Gut”: Their intestinal significance with and beyond butyrate, and prospective use as microbial therapeutics. Front. Microbiol. 13, (2023).

45. Gaudier, E. et al. Butyrate specifically modulates MUC gene expression in intestinal epithelial goblet cells deprived of glucose. Am. J. Physiol. Gastrointest. Liver Physiol. 287, 1168–1174 (2004).

46. Paone, P. & Cani, P. D. Mucus barrier, mucins and gut microbiota: The expected slimy partners? Gut 69, 2232–2243 (2020).

47. Vanhoutvin, S. A. L. W. et al. Butyrate-induced transcriptional changes in human colonic mucosa. PLoS One 4, (2009).

48. Kern, L., Kviatcovsky, D., He, Y. & Elinav, E. Impact of caloric restriction on the gut microbiota. Curr. Opin. Microbiol. 73, 102287 (2023).

49. Belzer, C. Nutritional strategies for mucosal health: the interplay between microbes and mucin glycans. Trends Microbiol. 30, 13–21 (2022).

50. Raba, G. & Luis, A. S. Mucin utilization by gut microbiota: recent advances on characterization of key enzymes. Essays Biochem. 67, 345–353 (2023).

51. Kwak, M. S., Cha, J. M., Shin, H. P., Jeon, J. W. & Yoon, J. Y. Development of a Novel Metagenomic Biomarker for Prediction of Upper Gastrointestinal Tract Involvement in Patients With Crohn’s Disease. Front. Microbiol. 11, 1–11 (2020).

52. Joossens, M. et al. Dysbiosis of the faecal microbiota in patients with Crohn’s disease and their unaffected relatives. Gut 60, 631–637 (2011).

53. Tailford, L. E., Crost, E. H., Kavanaugh, D. & Juge, N. Mucin glycan foraging in the human gut microbiome. Front. Genet. 6, 81 (2015).

54. Rajilić-Stojanović, M. et al. Global and deep molecular analysis of microbiota signatures in fecal samples from patients with irritable bowel syndrome. Gastroenterology 141, 1792–1801 (2011).

55. Xiao, L., Liu, Q., Luo, M. & Xiong, L. Gut Microbiota-Derived Metabolites in Irritable Bowel Syndrome. Front. Cell. Infect. Microbiol. 11, 1–11 (2021).

56. Glassner, K. L., Abraham, B. P. & Quigley, E. M. M. The microbiome and inflammatory bowel disease. Journal of Allergy and Clinical Immunology 145, 16–27 (2020).

57. Spiller, A. L. et al. Ultra-Processed Foods, Gut Microbiota, and Inflammatory Bowel Disease: A Critical Review of Emerging Evidence. Nutrients 17, 2677 (2025).

58. Belzer, C. et al. Microbial metabolic networks at the mucus layer lead to diet-independent butyrate and vitamin B12 production by intestinal symbionts. mBio 8, 1–14 (2017).

59. Donald, K. et al. Human milk IgA promotes normal immune development by limiting Th17-inducing Erysipelatoclostridium ramosum in the infant gut. Proceedings of the National Academy of Sciences 122, 2017 (2025).

60. Lawson, P. A., Perez, L. S. & Sankaranarayanan, K. Reclassification of Clostridium cocleatum, Clostridium ramosum, Clostridium spiroforme and Clostridium saccharogumia as Thomasclavelia cocleata gen. nov., comb. nov., Thomasclavelia ramosa comb. nov., gen. nov., Thomasclavelia spiroformis comb. nov. and T. Int. J. Syst. Evol. Microbiol. 73, 1–9 (2023).

61. Chia, L. W. et al. Deciphering the trophic interaction between Akkermansia muciniphila and the butyrogenic gut commensal Anaerostipes caccae using a metatranscriptomic approach. *Antonie van Leeuwenhoek*, International Journal of General and Molecular Microbiology 111, 859–873 (2018).

62. Fan, Y. et al. The gut microbiota contributes to the pathogenesis of anorexia nervosa in humans and mice. Nat. Microbiol. 8, 787–802 (2023).

63. Garcia, N. & Gutierrez, E. Anorexia nervosa and microbiota: systematic review and critical appraisal. Eating and Weight Disorders 28, (2023).

64. Marano, G. et al. Gut Microbiota in Women with Eating Disorders: A New Frontier in Pathophysiology and Treatment. Nutrients 17, 2316 (2025).

65. Pujo, J. et al. Bacteria-derived long chain fatty acid exhibits anti-inflammatory properties in colitis. Gut 70, 1088–1097 (2021).

66. Romaní-Pérez, M. et al. Holdemanella biformis improves glucose tolerance and regulates GLP-1 signaling in obese mice. FASEB Journal 35, 1–17 (2021).

67. Zhao, L. Y. et al. Role of the gut microbiota in anticancer therapy: from molecular mechanisms to clinical applications. Signal Transduct. Target. Ther. 8, (2023).

68. Brahe, L. K. et al. Specific gut microbiota features and metabolic markers in postmenopausal women with obesity. Nutr. Diabetes 5, e159–7 (2015).

69. Xu, T. et al. Berberis kansuensis extract alleviates type 2 diabetes in rats by regulating gut microbiota composition. J. Ethnopharmacol. 273, 2–10 (2021).

70. Min, Y. et al. Sex-specific association between gut microbiome and fat distribution. Nat. Commun. 10, 1–9 (2019).

71. Tropini, C. How the Physical Environment Shapes the Microbiota. mSystems 6, (2021).

72. Dey, P. & Ray Chaudhuri, S. The opportunistic nature of gut commensal microbiota. Crit. Rev. Microbiol. 49, 739–763 (2023).

73. Ducarmon, Q. R. et al. Remodelling of the intestinal ecosystem during caloric restriction and fasting. Trends Microbiol. xx, 1–13 (2023).

74. Peled, S. & Livney, Y. D. The role of dietary proteins and carbohydrates in gut microbiome composition and activity: A review. Food Hydrocoll. 120, 106911 (2021).

75. Louis, P. & Flint, H. J. Formation of propionate and butyrate by the human colonic microbiota. Environ. Microbiol. 19, 29–41 (2017).

76. Frolova, M. S., Suvorova, I. A., Iablokov, S. N., Petrov, S. N. & Rodionov, D. A. Genomic reconstruction of short-chain fatty acid production by the human gut microbiota. Front. Mol. Biosci. 9, 1–25 (2022).

## References

1. Reinhardt M, Thearle MS, Ibrahim M et al. A Human Thrifty Phenotype Associated With Less Weight Loss During Caloric Restriction. Diabetes 2015;64:2859–67.

