## Supplemental Data for "A 6-week all-liquid caloric restriction diet reduced abundance of some butyrate-producing bacteria while increasing abundance of mucin-degrading bacteria"

**Supplementary Material**

**Supplementary Table 1.**

**Table 1** General characteristics of the study participants [1]

| | **All (*n* = 12)** | **Men (*n* = 7)** | **Women (*n* = 5)** |
|---|---|---|---|
| Race, *n* | | | |
| Caucasian | 6 | 4 | 2 |
| Hispanic | 3 | 1 | 2 |
| Native American | 3 | 2 | 1 |
| Age (years) | 33.7 ± 8.6 | 32.4 ± 10.3 | 35.4 ± 6.1 |
| Height (cm) | 169.4 ± 10.8 | a | 162.0 ± 6.5 |
| Body weight (kg) | 107.5 ± 15.6 | 116.4 ± 8.1 | 95.1 ± 15.6* |
| Waist (cm) | 118.2 ± 18.0 | 121.0 ± 6.9 | 114.3 ± 8.5 |
| BMI (kg/$m^2$) | 38.0 ± 6.3 | 39.2 ± 6.8 | 36.2 ± 5.8 |
| Body fat (%) | 42.1 ± 8.2 | 36.6 ± 4.8 | 49.7 ± 4.9** |
| FM (kg) | 44.9 ± 9.5 | 42.9 ± 8.6 | 47.6 ± 11.0 |
| FFM (kg) | 62.6 ± 14.0 | 73.5 ± 2.2 | 47.5 ± 6.1** |
| 24h-EE (kcal/day) | 2,344 ± 363 | 2,591 ± 174 | 2,000 ± 250** |
| Daily intake during CR (kcal) | 1,536 ± 149 | 1,646 ± 79 | 1,404 ± 97** |
| 24-h respiratory quotient | 0.85 ± 0.04 | 0.85 ± 0.04 | 0.86 ± 0.03 |
| Fasting glucose (mg/dL) | 93.1 ± 5.0 | 92.0 ± 5.6 | 94.7 ± 3.9 |
| 2-h glucose (mg/dL) | 141.8 ± 39.1 | 142.0 ± 35.8 | 141.6 ± 47.7 |
| Fasting insulin (µIU/mL) | 19.7 ± 10.8 | 20.0 ± 6.0 | 19.3 ± 16.3 |
| 2-h insulin (µIU/mL) | 184.0 ± 172.9 | 172.4 ± 170.0 | 200.2 ± 195.6 |

Values are presented as mean ± SD unless otherwise indicated. Intake during CR are values from bomb calorimetry measurements. Differences between men and women were assessed using the Student *t* test: *$P < 0.05$; **$P < 0.01$.

**Supplementary Figure 1.**

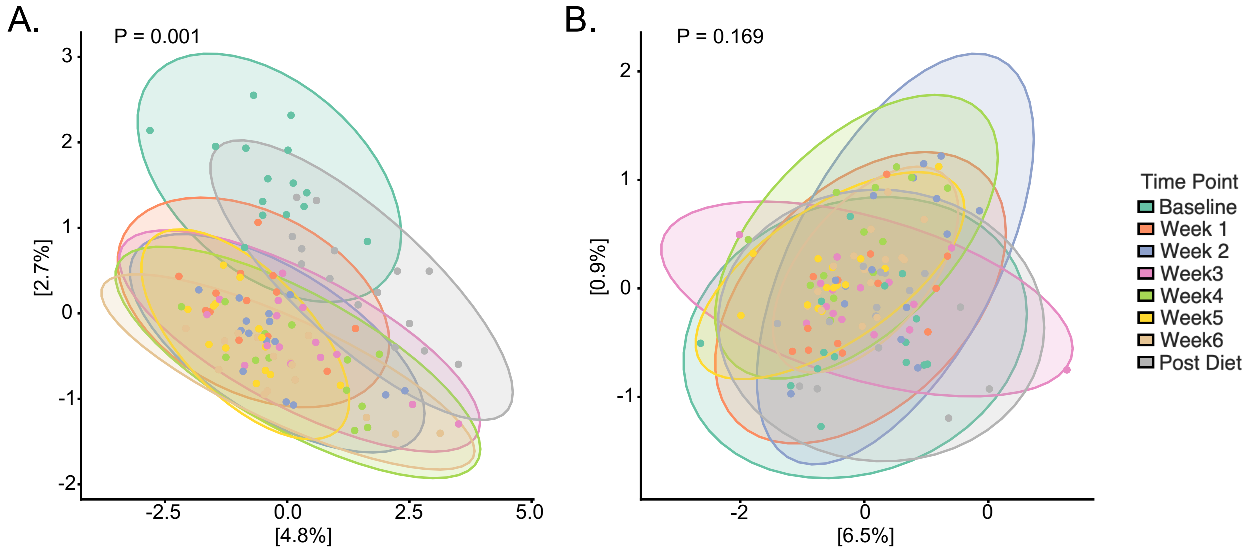


**Supplementary Figure 1. Unweighted and weighted Unifrac metrics for beta-diversity.** A) Unweighted unifrac was different by diet period (Permanova test, P = 0.001). B) Weighted unifrac was not different by diet period (Permanova test, P = 0.169).

### References


1. Reinhardt M, Thearle MS, Ibrahim M *et al.* A Human Thrifty Phenotype Associated With Less Weight Loss During Caloric Restriction. *Diabetes* 2015;**64**:2859–67.
